# Comparative gene expression analysis of differentiated terminal and lateral haustoria of the obligate root parasitic plant *Phelipanche ramosa* (Orobanchaceae)

**DOI:** 10.1101/2023.04.12.536555

**Authors:** Guillaume Brun, Julia K. H. Leman, Susann Wicke

## Abstract

Branched broomrape (*Phelipanche ramosa*, Orobanchaceae) is the most important parasitic weed of oilseed and vegetable crops in Europe. Deciphering the parasite’s basic biology and genetics underpinning adaptations to parasitism are essential for effective weed management. Here, we compared the gene expression profiles of differentiated terminal haustoria, formed immediately after the germination of *P. ramosa* with those of the parasite’s adventitious roots and lateral haustoria, which develop on the latter as it grows. Principal component analysis and functional annotation of differentially expressed genes indicate a greater transcriptional similarity between adventitious roots and lateral haustoria compared to terminal haustoria. Genes involved in hydrogen peroxide catabolic processes and mucilage metabolic processes were more abundant in lateral haustoria compared to adventitious roots, indicating that secondary host attachment involves secretion of adhesive compounds, cell wall modification, and a termination of the developmental program associated with root growth. In terminal haustoria, phytohormonal signaling pathways for post-germination growth and meristem development were the most prominent expression profile difference to lateral haustoria. Besides this, wide similarity of expressed genes between lateral and terminal haustoria suggest overlapping pathways underlying haustorium differentiation and a conserved functional relevance of both haustoria types at maturity. Together, our study provides first insights into the transcriptional landscapes of the primary organs securing *P*.*ramosa*’s nutritional success. However, more research is needed to investigate whether the parasite’s various haustorium types differ in their sensitivities to environmental stimuli and whether transcriptional divergence in terminal and lateral haustoria of *P. ramosa* reflect differences regarding their developmental origin.

## Introduction

*Phelipanche ramosa* (Orobanchaceae) is a noxious parasitic weed that threatens various host plant species belonging to Fabaceae, Solanaceae, Brassicaceae, and Asteraceae, from the Middle East to North America (Parker, 2013). It causes losses of tens of millions of euros in annual output and control initiatives. Completely non-photosynthetic, it strictly requires a host to fulfill its lifecycle. Seeds must perceive host-derived compounds to germinate and differentiate a terminal haustorium, a specialized and multifunctional organ, which attaches and penetrates a host root, and then connects to the host vascular system to withdraw the water and nutrients necessary for its growth (Brun *et al*., 2021). Subsequent maturation of the parasite involves the differentiation of a tubercle that develops adventitious roots as resources accumulate.

Adventitious roots of *P. ramosa* may branch and produce lateral haustoria upon contact to host roots. Anatomical work suggests that undifferentiated parasite cells develop a labyrinthine wall adjacent to host xylem before establishing a functional connection between the lateral haustorium and a host root (Dörr & Kollmann, 1975). Those lateral haustoria of *P. ramosa* form no direct phloem connection, but rather a phloic conduit consisting of two types of parenchyma cells, with contact cells bridging the host sieve tubes and transition sieve cells interconnected by numerous plasmodesmata (Dörr & Kollmann, 1975). In line with this, the presence of phloem cells in the neck of mature lateral haustoria have rarely been observed, in contrast to that of the holoparasite’s terminal haustoria (Joel, 2013).

Knowledge beyond structural aspects of lateral haustoria in broomrapes is scarce. For example, it remains unclear if *P. ramosa*’s lateral haustoriogenesis also depends on host-derived chemicals. Along this line, also the developmental programs triggering and underlying the differentiation of adventitious roots and lateral haustoria are unknown. Here, we aim to gain a first understanding as to whether lateral haustoria of holoparasites such as broomrape have similar or unique functions when compared to the terminal haustoria. We tackle this research question using a comparative transcriptomic approach to explore the transcriptional landscapes of matured terminal haustoria, adventitious roots, and lateral haustoria using *P. ramosa* as a representative model.

### Material & Methods

#### Plant material

*Phelipanche ramosa* seeds were sterilized and conditioned for twelve days as previously described (Brun *et al*., 2019), and applied to *rac*-GR24-treated semi-*in vitro* grown roots of 14d-old tomato seedlings (cv. ‘Zuckertraube’). To obtain material of differentiated and functional primary attachments, we collected six biological replicates of terminal haustoria seven to ten days after infestation. Between ten to twelve weeks after infestation, we collected branching, non-haustorialized adventitious roots from parasite tubercles, and a triplicate of fully developed lateral haustoria was sampled from adventitious roots ca. seven to twelve days, i.e., after host penetration was evident. Adventitious root samples as well as terminal and lateral haustoria samples were carefully severed from the host root and adjacent own tubercle tissue. All samples were immediately snap-frozen in liquid nitrogen and stored at - 80° C until further use.

#### Sample preparation for RNA sequencing

Total RNA was extracted using the PicoPure™ RNA Isolation Kit (ThermoFisher Scientific) with on-column RNAse-free DNAse treatment (Qiagen) following manufacturer’s instructions. RNA quality and quantity were assessed using an Agilent 2100 Bioanalyzer Instrument and a Qubit4 fluorometer, respectively. RNA sequencing was carried out by Eurofins Genomics (Konstanz, Germany), where poly-A capture libraries were prepared from >0.1 µg of total RNA using the TruSeq Stranded mRNA Library Prep kit (Illumina). Sequencing was performed on an Illumina NovaSeq 6000 platform in 150 bp paired-end mode, targeting 30 million read pairs per library.

#### Transcriptomics analyses

Adaptors and low-quality paired-end sequences were removed from the raw reads using Trimmomatic v0.36 (Bolger *et al*., 2014), after which read quality was assessed using FastQC v0.11.9 (Andrews, 2010). We used Bowtie2 v2.4.1 (Langmead & Salzberg, 2012) to map trimmed reads to the OrAeBC5 reference build from the PPGP database (http://ppgp.huck.psu.edu), which we reannotated using the Trinotate pipeline (https://github.com/Trinotate/Trinotate/wiki); read counts were estimated using RSEM v1.3.3 (Li & Dewey, 2011). Differential gene expression analyses across all pairwise contrasts were carried out using the DESeq2 package (Love *et al*., 2014). Genes were considered differentially expressed at a log2 fold-change threshold of 2 and an adjusted *P-* value (FDR) threshold of 0.01. Gene Ontology (GO) enrichment analyses were conducted using enricher function of the ‘clusterProfileR’ package (Wu *et al*., 2021).

## Results

To obtain first insight into the transcriptional landscape of *P. ramosa’s* nutritionally relevant organs, we compared the gene expression profiles of its adventitious roots with its fully differentiated terminal and lateral haustoria (Fig. 1A). Principal component analysis of regularized log-transformed RNAseq count data highlighted high dissimilarity in transcript abundance between terminal haustoria and the two other tissue types, while much lower variance was observed between lateral haustoria and adventitious roots (Fig. 1B). Pairwise comparisons revealed 406 differentially expressed genes (DEGs) between adventitious roots and lateral haustoria, as opposed to 3047 and 3288 DEGs when comparing adventitious roots or lateral haustoria to terminal haustoria, respectively. Nevertheless, the actual total of DEGs was 3925 across all pairwise comparisons, which could be explained by the extensive overlap in the identity of the DEGs when comparing adventitious roots or lateral haustoria to terminal haustoria (Fig. 1C). We identified two main expression groups, whereby most of the highly expressed genes in terminal haustoria were lowly expressed in adventitious roots and lateral haustoria, and vice versa (Fig. 1D).

**Figure 1.**
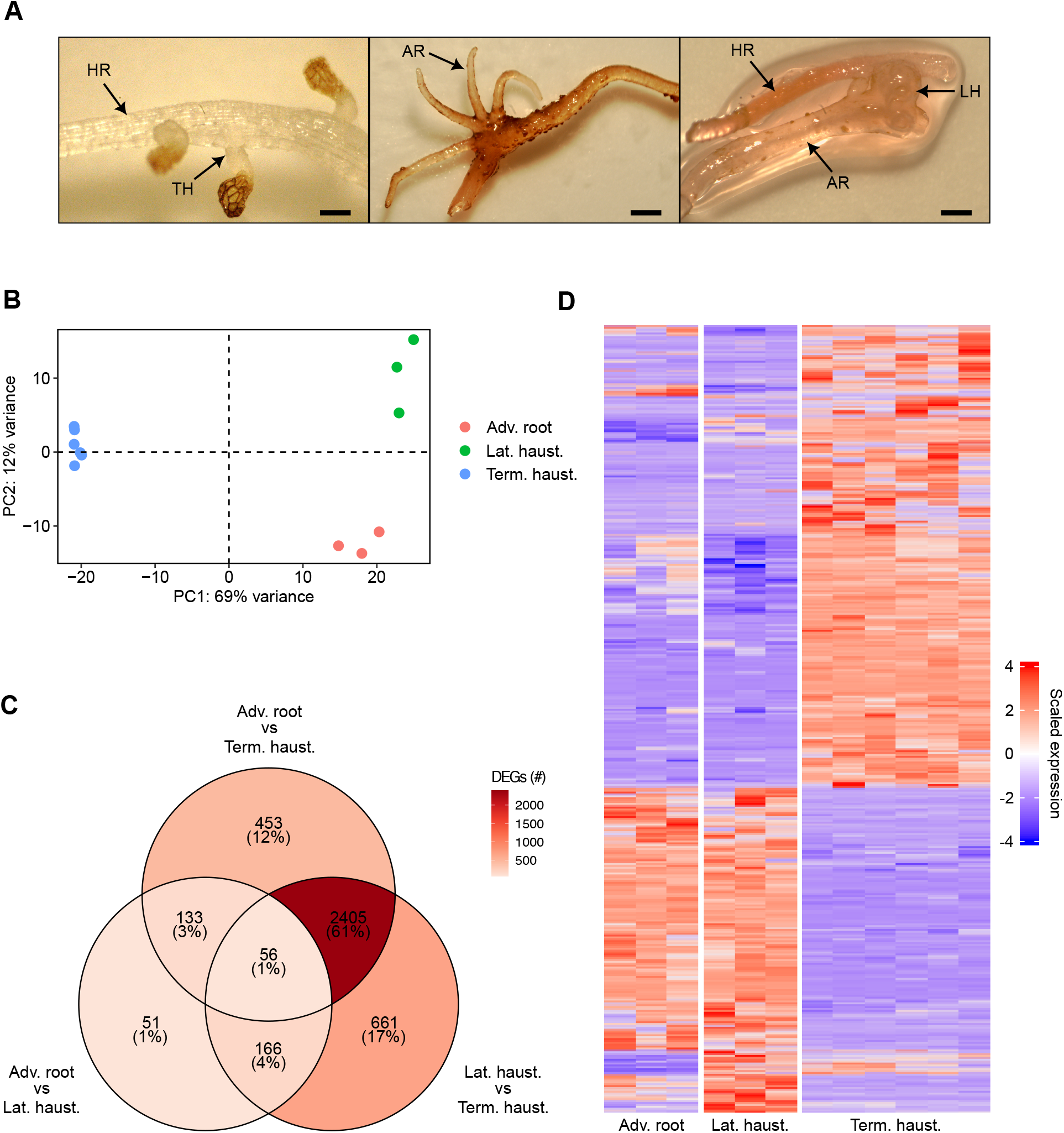
Phenotypic and gene-level expression differences between terminal haustoria, lateral haustoria, and adventitious roots in *Phelipanche ramosa*. **(A)** Representative developmental stages (scale = 200 µm). HR, Host root; TH, Terminal haustorium; AR, Adventitious root; LH, Lateral haustorium. **(B)** Biplot of the first and second principal components (PC1 vs. PC2) generated from regularized log-transformed count data of each sequencing dataset. **(C)** Venn Diagram of the number of significant differentially expressed genes across all pairwise comparisons (log2 fold-change > 2 and adjusted *P*-value < 0.01). **(D)** Clustered heatmap generated from regularized log-transformed count data of all significant differentially expressed genes across all datasets.

Detailed inspection of the extent and direction of differential expression revealed a fairly equal share of up-regulated and down-regulated genes within each comparison (Fig. 2A).

**Figure 2.**
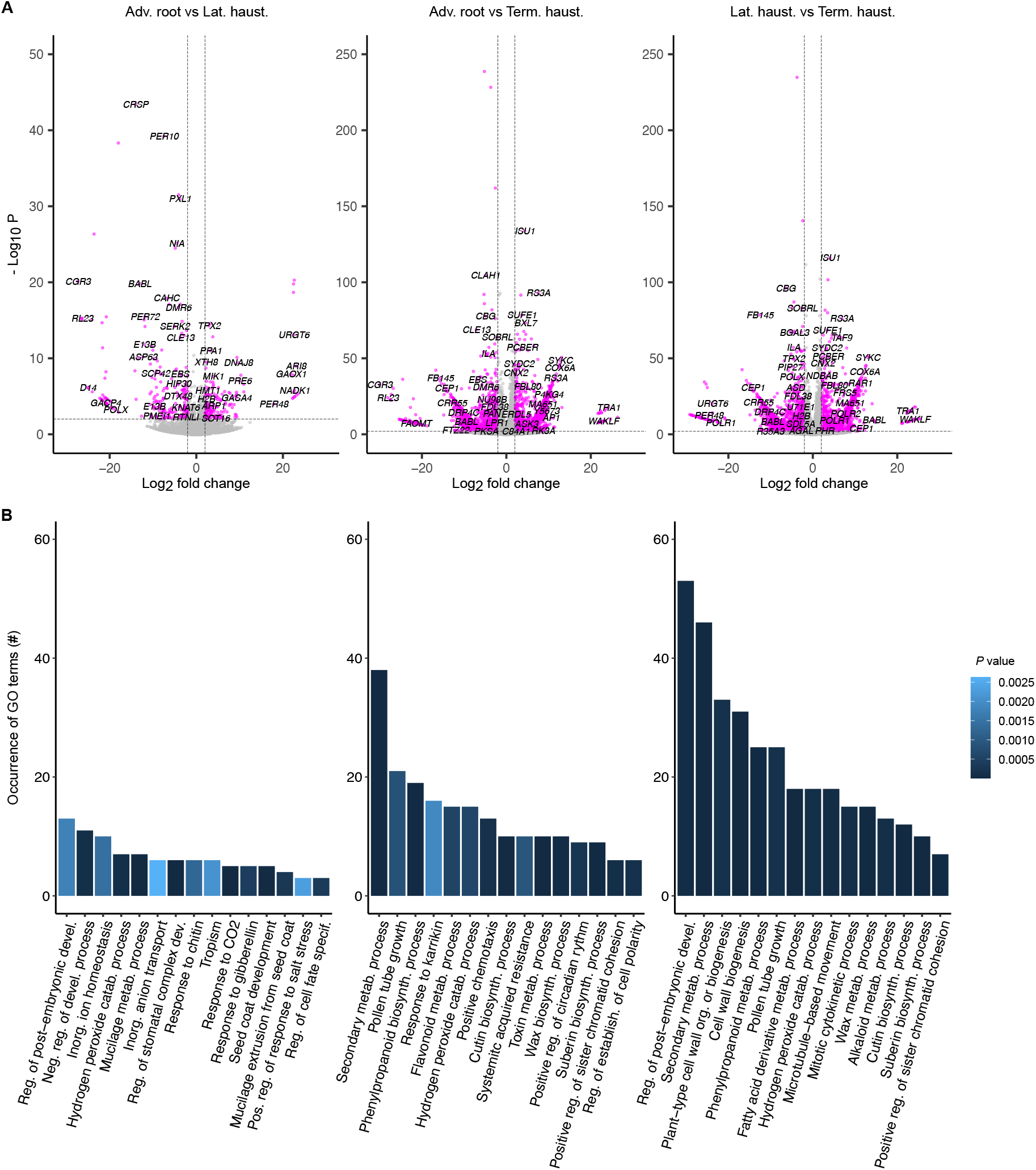
Gene-level comparisons of expression and functional annotation between terminal haustoria, lateral haustoria, and adventitious roots in *Phelipanche ramosa*. **(A)** Volcano plots displaying the distribution of significant differentially expressed genes across all pairwise comparisons (magenta dots; log2 fold-change > 2 and adjusted *P*-value < 0.01). Labels correspond to a subset of gene names of homologous sequences within the Viridiplantae family. Y-axes limit to the lower adjusted *P-*value for each dataset in negative Log10 mode. Positive and negative log2 fold change values indicate significant gene up-regulation and down-regulation in the left term, respectively. **(B)** Overview of the 15 most significant abbreviated GO terms belonging to the ‘biological process’ category across all pairwise comparisons. Ties with redundant parent or child terms were manually excluded from the plots, but are accessible in Table S1.

Functional annotation of the OrAeBC5 reference build enabled to annotate 2119 of all DEGs (53.98%), 1864 of which having been assigned to a homologous sequence belonging to the Viridiplantae family. GO enrichment analyses revealed a large proportion of overlapping GO terms when comparing terminal haustoria to either of the two other tissue types (Fig. 2B, Fig. S1). Of the 165 significantly enriched terms within the ‘biological process’ category that were found when comparing adventitious roots to lateral haustoria, 97 DEGs (58.8%) did not intersect with any other comparison (Fig. S1A). This data suggests a higher functional similarity between adventitious roots and lateral haustoria compared to terminal haustoria.

Enrichment analyses revealed that most of the significantly enriched GO terms encompass genes that were generally more highly expressed in lateral haustoria compared to adventitious roots (Fig. 2B). For instance, the category ‘hydrogen peroxide catabolic process’ contains seven DEGs annotated as peroxidases (*PER10, PER48, PER52*, and *PER72*), of which only *PER48* is more abundant in adventitious roots. This also applies to six of the seven DEGs associated to ‘mucilage metabolic process’, which encode homologs of the subtilisin-like proteases SBT1.7 and SBT1.8, a putative glycosyltransferase GT7, and the trihelix transcription factor GTL1. Subtilisin-like proteases and glycosyltransferases are involved in biosynthesis of mucilage backbones, xylan and pectin, respectively (Rautengarten *et al*., 2008; Phan *et al*., 2016), while GTL1 negatively regulates root hair growth (Shibata *et al*., 2022). These findings suggest that attachment of lateral haustoria to host roots is facilitated by the secretion of adhesive compounds and cell wall components and followed by a termination of the developmental program associated with root growth. However, it is rare that most of a gene set associated to a given GO term is more highly expressed in one tissue compared to another. For example, the term ‘tropism’ contains three genes encoding homologs of the kinase D6PKL3, the E3 ubiquitin ligase WAV3, and the microtubule-associated protein EB1B that are more expressed in adventitious roots, whereas three genes encoding homologs to the heat shock transcription factor HSFA4C, the signaling peptide CEP3, and the auxin efflux carrier PIN2 are more expressed in lateral haustoria. The latter finding is reminiscent to the pivotal role for auxin synthesis in differentiation of lateral haustoria in the facultative hemiparasite *Phtheirospermum japonicum* (Ishida *et al*., 2016).

Six of the 13 DEGs associated with ‘positive chemotaxis’ were more expressed in terminal haustoria compared to adventitious roots, which includes those encoding the SNF1-related protein PV42A, the Ser/Thr-protein kinase SIK1, the eLRR kinase PXL1, the receptor kinase MIK2, and the nucleoredoxins NRX1 and NRX11. The seven other genes involved in positive chemotaxis were more expressed in adventitious roots, included three homologs of the nucleoredoxin NRX1, the inositol polyphosphate kinase IPK2B, the LRR extensin-like LRX7, the Ser/Thr-protein kinase PBL24, and the receptor-like cytoplasmic kinase LIP1. Except for genes involved in secondary metabolic processes, toxin metabolism, and a positive regulation of circadian rhythm, the most significantly enriched terms when comparing adventitious roots to terminal haustoria revealed genes that are overall more expressed in terminal haustoria. This includes gene sets involved in suberin, cutin, and wax biosynthesis, such as genes encoding the alkane hydroxylase CYP96A15, the E3 ubiquitin ligases CER9, the aldehyde decarbonylase CER1, and the acyl-coA reductase CER4. These same three sets are also globally more expressed in terminal haustoria compared to lateral haustoria, as are all most significantly enriched categories except for the ‘secondary metabolic process’ (Fig. 2B).

The most highly enriched category between lateral and terminal haustoria was the ‘regulation of post-embryonic development’, among which 35 of the 53 corresponding genes were more expressed in terminal haustoria. This includes genes involved in post-germination growth and meristem development within the cytokinin, brassinosteroid, abscisic acid, gibberellin and ethylene signaling pathways. Inversely, of the genes more expressed in lateral haustoria we could find those encoding HD3A, the rice ortholog to the *Arabidopsis* FT protein, and the trihelix transcription factor PTL, both involved in flower development.

## Discussion

This study shows that only a few transcriptional changes occur after the differentiation of lateral haustoria on adventitious roots of *P. ramosa*. Most genes that were highly expressed in adventitious roots and lateral haustoria were present in low abundance in terminal haustoria, and vice versa. The current model proposes that parasitic plants secrete peroxidases and H2O2 from radicle tips, which enable to convert host-released phenolic acids into quinones to initiate haustorium differentiation (Yoshida *et al*., 2016). The herein found high levels of expression of peroxidases in both lateral and terminal haustoria suggests overlapping pathways underlying haustorium differentiation. In line with this, most genes associated with mucilage synthesis were equally more abundant in lateral and terminal haustoria, implying similarities in host attachment systems and the formation of haustorial plates.

It remains uncertain whether the earliest phases of haustorium induction and the actual vascular connection processes are similar between broomrape’s terminal and lateral haustoria. After all, both types of broomrape haustoria differ cellularly regarding their vascular connections (Dörr & Kollmann, 1975). Cell identity and origin at the parasite-host interface are more clearly distinguishable at lateral attachment points while parasite and host cells mingle and fuse more strongly at the terminal haustorial attachment (Dörr & Kollmann, 1995).

We found different protein families, mostly falling within the kinase superfamily, associated to chemotaxis in lateral and terminal haustoria, yet Ser/Thr-protein kinases and neoredoxins (thioredoxin-disulfide reductase activity) were notably overrepresented in lateral haustoria. It might be that diverging phosphorylation and redox signaling pathways, hence, perhaps diverging host signals, are required for the radicle and the root tips to grow towards the host roots. According to the analysis of the “secondary metabolic process” category, the majority of genes, whether expressed more in terminal or lateral haustoria, appeared to be closely connected to lignin and suberin biosynthesis.

More research is required to determine whether the two types of haustoria have different lignin composition patterns. It is also unknown whether the two types of haustoria exhibit contrasted sensitivities to different host lignins given that host-derived lignins also act as a matrix in the synthesis of HIFs (Yoshida *et al*., 2016). In addition, the higher levels of gene transcription associated with post-embryonic growth in terminal haustoria suggest that the developmental programs associated to terminal haustoriogenesis overlap largely with seed-related pathways (Holdsworth *et al*., 2008). This scenario is sensible given that terminal haustoriogenesis is consequential to seed germination within a very short time span. Repression of embryonic properties during seed germination of non-parasitic plants ensures proper seedling establishment (Tanaka *et al*., 2008). We, therefore, expect interdependency between seed germination and terminal haustoria signaling pathways in order to guarantee the proper establishment of the primary connection to the host and subsequent early parasite development. The divergence of transcriptional landscapes in terminal and lateral haustoria of *P. ramosa* may also reflect differences regarding their developmental origin. Terminal haustoria likely develop from root apical meristem-like cells clusters in an embryonic root, whereas lateral haustoria probably develop from re-meristematized cortical cells of a fully differentiated adventitious root (Attawi & Weber, 1980). The exact origins of the various *P. ramosa* haustoria remains unclear to date. Whether or not the earliest time points of terminal and lateral haustorium induction truly underly the observed transcriptional departures between the two haustorium type of broomrape or whether the herein described differences occur in conjunction with functional specificity deserve more attention in the future.

## ACKNOWLEDGMENTS

We thank the Delavault lab at the University of Nantes (France) for sharing with us with seeds of *Phelipanche ramosa*. Thanks are also due to Alina Griese for technical and horticultural assistance. This research received financial support from Deutsche Forschungsgemeinschaft (DFG grant WI4507/3-1 to S.W.), which is gratefully acknowledged. J.K.H.L. is a fellow of the Elsa Naumann graduate program of the state of Berlin, Germany.

## Author Contributions

Research designed: J.K.H.L and S.W.; Experiments: J.K.H.L; Data analysis: G.B., J.K.H.L., and S.W.; Manuscript writing: G.B., J.K.H.L., and S.W.

## Data availability

All RNASeq data-related files, incl. read count tables and analysis scripts, are available for download from Mendeley Data, doi: 10.17632/ rjrbd6sjdy.1 and the NCBI Short Read Archive.

## Supplemental Material

**Supplemental Figure 1** Overrepresentation analysis of differentially expressed genes.

**Supplemental Table 1** Lists of enriched GO terms from pairwise comparisons (*P* value < 0.05)

