## Supplementary material for "Comparative gene expression analysis of differentiated terminal and lateral haustoria of the obligate root parasitic plant *Phelipanche ramosa* (Orobanchaceae)": Fig. S1

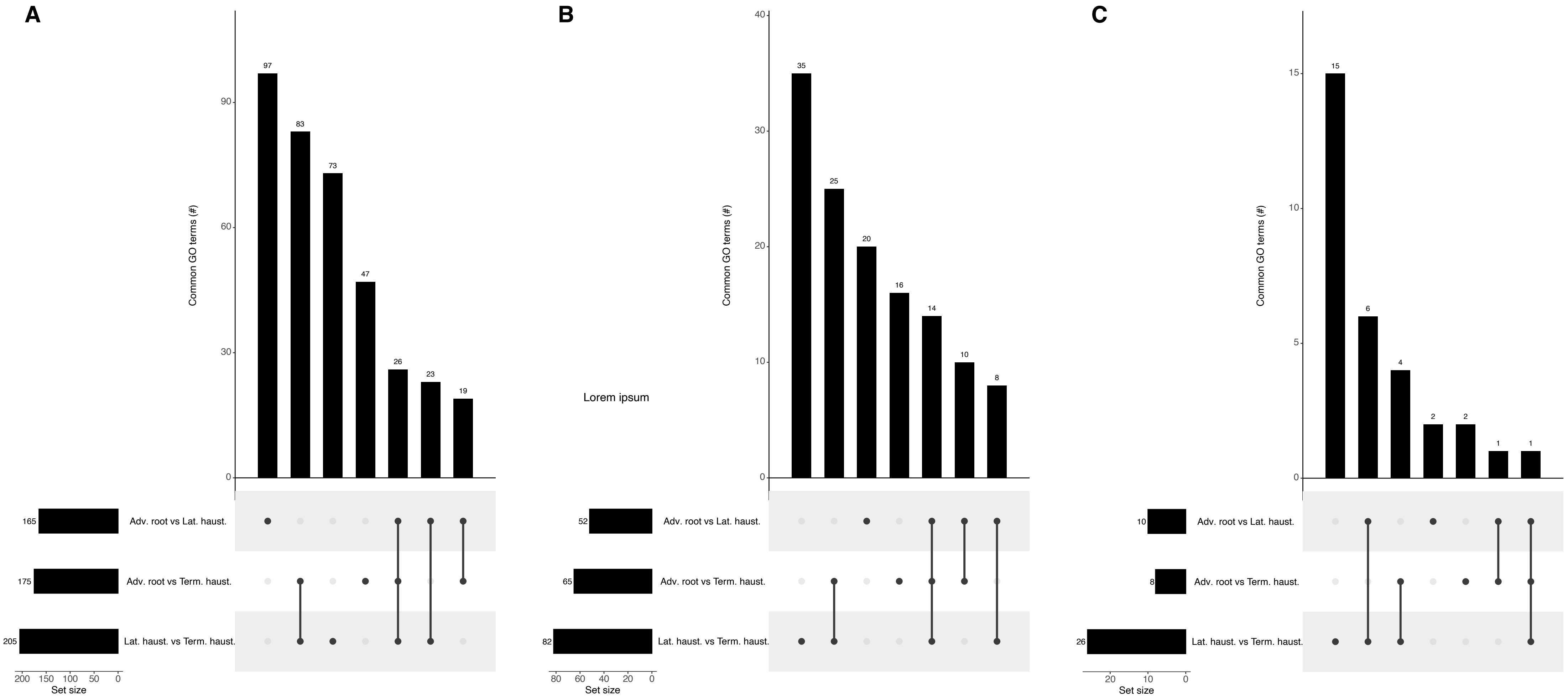


**Supplemental Figure 1 Overrepresentation analysis of differentially expressed genes.** Upset plots summarizing the intersections between significantly enriched GO terms belonging to **(A)** 'biological process', **(B)** 'molecular function', and **(C)** 'cellular component' catego-ries across all pairwise comparisons (P value < 0.05).
